# Rapid HIV-1 capsid interaction screening using fluorescence fluctuation spectroscopy

**DOI:** 10.1101/2020.11.13.382242

**Authors:** Derrick Lau, James C. Walsh, Claire F. Dickson, Andrew Tuckwell, Jeffrey H. Stear, Dominic J. B. Hunter, Akshay Bhumkar, Vaibhav Shah, Stuart G. Turville, Emma Sierecki, Yann Gambin, Till Böcking, David A. Jacques

## Abstract

The HIV capsid is a multifunctional protein capsule for delivery of the viral genetic material into the nucleus of the target cell. Host cell proteins bind to a number of repeating binding sites on the capsid to regulate steps in the replication cycle. Here we develop a fluorescence fluctuation spectroscopy method using self-assembled capsid particles as the bait to screen for fluorescence-labelled capsid-binding analytes (‘prey’ molecules) in solution. The assay capitalizes on the property of the HIV capsid as a multivalent interaction platform, facilitating high sensitivity detection of multiple prey molecules that have accumulated onto capsids as spikes in fluorescence intensity traces. By using a scanning stage, we reduced the measurement time to 10 s without compromising on sensitivity, providing a rapid binding assay for screening libraries of potential capsid interactors. The assay can also identify interfaces for host molecule binding by using capsids with defects in known interaction interfaces. Two-color coincidence detection using fluorescent capsid as bait further allows quantification of binding levels and determination of binding affinities. Overall, the assay provides new tools for discovery and characterization of molecules used by HIV capsid to orchestrate infection. The measurement principle can be extended for the development of sensitive interaction assays utilizing natural or synthetic multivalent scaffolds as analyte-binding platforms.

## INTRODUCTION

Early in infection, HIV releases its capsid containing genomic RNA and associated proteins into the cytoplasm of the target cell. The HIV capsid is a pleomorphic protein nanocontainer with a length of ~120 nm and a width of ~60 nm. It is assembled from ~1500 capsid protein (CA) monomers that form a cone- or pill-shaped lattice consisting of mainly hexamers (~250) and exactly 12 pentamers – a geometric constraint required for surface closure.^1^ The capsid is involved in many processes of the viral replication cycle. It serves as a reaction container for efficient conversion of the HIV genomic RNA into DNA^2,3^ while protecting its genetic material against degradation or detection by cellular sensors that would elicit innate immune responses.^4^ The capsid also functions as a vehicle for transport across the cytoplasm,^5–7^ import into the nucleus^8^ and site selection for integration of the viral DNA into host cell chromosomes.^9^

To regulate these processes, the viral capsid has evolved to bind proteins and small molecules from the host cell at different stages of the replication cycle after cell entry. The host cell also contains molecules of the innate immune system that compete for binding to the capsid and reroute the virus to degradative pathways. These host molecules bind to different repeating interfaces on the capsid that are located either on each CA subunit or between CA subunits within or between hexamers of the lattice.

As the capsid must orchestrate so many processes, it is likely that a substantial number of cofactors (used by the virus to infect) and restriction factors (used by the cell to defend against the virus) remain to be discovered and characterized. Assembling a comprehensive list of capsid binders requires biochemical and biophysical methods for detection and quantification of the complexes formed between HIV capsid and various analytes *in vitro*. Since authentic HIV capsids are unstable and difficult to isolate, many approaches utilize stabilized structures of engineered CA that self-assemble into defined parts of the capsid lattice, such as crosslinked CA hexamers and pentamers^10^ or the recently developed suite of crosslinked CA assemblies ranging from 2–42 CA molecules for identifying specific cofactor binding sites on the capsid lattice.^7,11^ Binding of analytes to these soluble capsid assemblies has been measured qualitatively using label-free assays such as differential scanning fluorimetry ^2,3,12^ and gel filtration^7,11^ or quantitatively using isothermal titration calorimetry^13,14^ and surface plasmon resonance spectroscopy.^15,16^ Alternatively, binding to capsid can be detected using spin-down assays with larger CA assemblies such as crosslinked CA tubes^17^ or isolated viral cores,^18–21^ whereby protein analytes that co-pellet with CA tubes (or viral cores) are detected by Western blotting. Recently, we also described fluorescence biosensors to quantify binding of fluorescence-labelled analytes to surface-immobilized CA tubes or spheres.^22,23^

Here we develop an analytical method to facilitate rapid screening of fluorescence-labelled analyte binding to capsid in solution, a capability that complements existing approaches. The method builds on single molecule spectroscopy approaches for detection of protein oligomerization and aggregation^24–26^ that monitor in real time the light emitted from fluorescence-tagged protein assemblies diffusing through a confocal excitation volume (~1 fL).^27^ Formation of analyte-capsid complexes is detected by analysis of the analyte’s fluorescence intensity traces using a method based on Number and Brightness,^28^ or by two-color coincidence detection (TCCD)^29^ in measurements with labelled analyte and labelled capsid. Trace acquisition is accelerated by using a scanning stage, which enables rapid detection of multiple events despite the slow diffusion of the large capsid particles. Overall, the approach is suited for rapid discovery of new capsid binders and identification of interaction sites.

## MATERIALS AND METHODS

### Expression, purification and labelling of proteins

HIV-1 CA proteins (K158C, A204C, R18G/A204C, N57D/A204C and N74D/A204C) were expressed in bacteria and purified as previously described.^23^ CA K158C was labelled with Alexa Fluor 568-C5-maleimide (AF568, Thermo Fisher Scientific, A20341) as previously described.^23^ Untagged recombinant CypA was expressed, purified and labelled with Alexa Fluor 488-C5 maleimide (AF488, Thermo Fisher Scientific, A10254) as previously described.^16^

### Labelling of peptides

CPSF6 peptide (P313-G327) with C-terminal cysteine (PVLFPGQPFGQPPLGC, Genscript) and FEZ1 peptide (M174-L188, with N-terminal extension containing a cysteine residue (KCGGSGGMMQNSPDPEEEEEVL, Genscript) were labelled with Alexa Fluor 488-C5-maleimide at 1:1 molar ratio in 50 mM Tris, pH 8 and quenched with 1 mM DTT. The labelled peptides (designated as CPSF6-AF488 and FEZ1-AF488) were snap frozen in liquid nitrogen and stored at −40 °C.

### Labelling of His_6_-tag proteins with tris-NTA-AF488

A 200 μM aqueous solution of tris-NTA-AF488 was mixed with an equal volume of 1 mM aqueous NiCl_2_ to prepare Ni-loaded dye. His_6_-CypA or His_6_-LcrV was diluted to a concentration of 33 μM with 50 mM Tris pH 8 and Ni-loaded dye was added at a 1:1.5 molar ratio. Unbound dye was removed using a Zeba spin column (Thermo Fisher Scientific, 89882). The concentration of labelled His_6_-proteins was determined using the absorbance at 495 nm and the extinction coefficient of tris-NTA-AF488 (ε_495 nm_ = 71000 M^−1^cm^−1^).

### Cell-free protein expression

The coding sequences for CypA, CPSF6 (isoform 2) and a fragment of NUP153 (residues 1407-1423) were cloned into the plasmid pCellFree_03 for expression with N-terminal GFP tag.^30^ Each plasmid (60 nM) was added to *Leishmania tarentolae* extract^31,32^ supplemented with RnaseOUT (1:1000, Invitrogen) and the mixture was incubated at 28 °C for 2.5 h.

### Assembly of HIV-1 capsid particles

To initiate assembly, CA A204C was diluted to 80 μM in 50 mM Tris, pH 8, 1 M NaCl and incubated at 37 °C for 15 min and then at 4 °C overnight. Labelled capsids were produced by co-assembly of CA A204C and CA K158C-AF568 (76 μM:4 μM) as above, yielding particles with an estimated label fraction of 2.5% due to the lower assembly efficiency of labeled CA K158C.^22,23^ For affinity measurements, assembled particles were collected by centrifugation (18000 g, 10 min, 4°C) and resuspended in 50 mM Tris, pH 8, 1M NaCl to remove unassembled CA.

### Negative staining electron microscopy

Capsid assembly reactions (5 μL) were applied to a Formvar/carbon Cu grid (Ted Pella, 01811), briefly incubated and wicked dry. The sample was stained with uranyl acetate solution (2% w/v) and immediately wicked dry. This step was repeated twice, and the grid was air dried. Transmission electron micrographs were collected using the FEI Tecnai G2 20.

### Brightness analysis and TCCD

Fluorescence traces of CypA-AF488 (50 nM), CPSF6-AF488 (250 nM), FEZ1-AF488 (100 nM) were measured in the presence or absence of capsid particles (final concentration equivalent to 12 μM monomeric CA) in 50 mM Tris, pH 8; 150 mM NaCl; 5 mM MgCl2; 0.1% w/v BSA. 20 μL samples were added to a custom polydimethylsiloxane plate adhered to a glass coverslip (ProSciTech, G425-7080) and observed on an inverted microscope equipped with 488 nm (2.6 mW) and 561 nm (0.5 mW) lasers and a water immersion 40x/1.2 NA objective (Zeiss). Emitted fluorescence was separated into two channels using a dichroic mirror (565 nm) and filtered through a 525/50 nm band pass filter (AF488/GFP signal) and 590 nm long pass filter (AF568 signal), respectively, prior to focusing onto separate single photon avalanche diodes (Micro Photon Devices). Fluorescence traces were recorded for 100 s (10 s/trace) in 1 ms bins using a scanning stage operated at 1 μm/s. All binding experiments using recombinant proteins had at least three independent repeats. Cell-free expressed N-terminal GFP fusion protein levels were adjusted to ~2500 photon count (~50 nM) for binding experiments. Inhibitors included cyclosporin A (CsA, 5 μM), hexacarboxybenzene (HCB, 10 μM) and CPSF6 peptide (100 μM, PVLFPGQPFGQPPLG, Genscript).

### Analysis of fluorescence traces

Fluorescence traces were analyzed using the custom software TRISTAN (Two Reagents Incident Spectroscopic Analysis, freely available on https://github.com/lilbutsa/Tristan). Traces were filtered to exclude signals outside the linear range of the detectors.

## RESULTS AND DISCUSSION

### Fluorescence fluctuation spectroscopy assays for the detection of HIV-1 capsid-binding cofactors

Fluorescence fluctuation spectroscopy relies on detecting photons emitted by fluorescent molecules freely diffusing in and out of the confocal volume of a microscope (Fig 1A). Here we adapted this technique to detect the binding to capsid particles of a range of fluorophore-labelled host proteins and small molecules known to function as cofactors for HIV infection (Fig 1A, inset). We prepared HIV-1 capsid particles by self-assembly of recombinant CA A204C^33^ at high salt concentrations (1 M NaCl), resulting in the formation of conical and tubular lattices with capped ends as seen by negative staining electron microscopy (Fig 1B). These lattices resemble authentic HIV capsids in that they are primarily hexameric and include pentamers at the end caps to achieve lattice closure. The engineered cysteine residues form disulfide linkages that prevent lattice disassembly at physiological salt concentrations.^33^ For two-color experiments, capsid particles were rendered fluorescent by incorporating a small fraction of the AF568-labelled CA K158C into the CA A204C lattice during self-assembly.^23^

**Fig 1.**
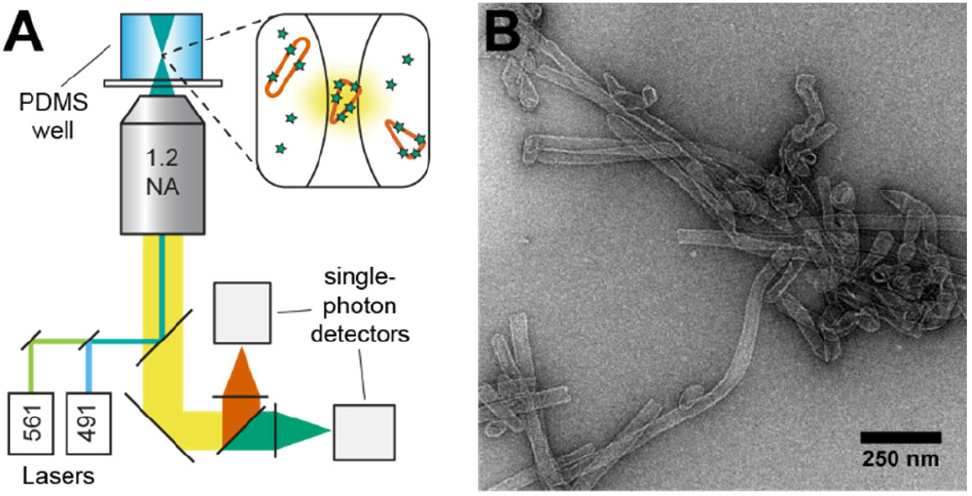
Fluorescence fluctuation spectroscopy assay for detection of capsid binders. **A.** Schematic of the microscope set-up. Inset: Fluorescence emission from a complex formed between analyte molecules (green stars) and CA A204C cones or tubes (vermillion) moving through the detection volume. **B.** Negative staining electron micrograph of disulfide cross-linked tubes and cones formed by self-assembly of CA A204C for use as the bait.

The binding assay was carried out by adding a mixture of capsid particles and labelled analyte into a well of a 192-well polydimethylsiloxane device adhered to a glass coverslip. We then measured fluorescence intensity (photon counting) traces as a function of time using an inverted confocal microscope with custom-built two-color excitation and two-channel single-molecule detection modules (Fig 1A). During acquisition, the sample stage was displaced at a constant speed such that the confocal volume “scans” the well to increase the rate of detecting the slowly diffusing capsid particles. Binding of fluorescent analyte (‘prey’) to capsid particles (‘bait’) was detected in two modes: (1) via changes in the brightness parameter of the analyte signal recorded in the absence and presence of capsid particles (single-color measurements) or (2) via coincidence analysis of intensity peaks in analyte and capsid traces (two-color measurements). Below we characterize these modes using analytes labelled with chemical fluorophores or labelled by fusion to a fluorescent protein.

### Brightness analysis detects accumulation of fluorescent analytes onto capsid particles

Single-color detection of capsid-binding fluorescent analytes is based on the Number and Brightness method^28^ and relies on differences in the analyte intensity fluctuations measured in the absence and presence of capsid (Fig 2A). We first tested this approach using chemically labelled HIV cofactors that are known to bind to capsid after cell entry, namely AF488-labelled CypA,^34,35^ an AF488-labelled peptide derived from CPSF6 (residues 313-327)^14^ and fluorescein-labelled dATP.^3^ Brightness measurements were carried out at analyte concentrations in the mid-high nM range. Under these conditions, the average number of (monomeric) molecules in the confocal volume exceeds 10 molecules/ms,^36^ giving rise to background intensity traces characterized by small fluctuations around the mean (Fig 2B, top). In contrast, co-diffusion through the confocal volume of multiple fluorescent analytes bound at equilibrium onto a capsid particle leads to the appearance of large intensity spikes (Fig 2B, middle). We note that a conical capsid particle contains about 1500 binding sites for CypA and CPSF6 peptide and 250 binding sites for dATP;^37^ tubular particles and/or small aggregates of several particles would present a greater surface area and hence larger number of analyte binding sites. As a result of the large number of analytes that can be bound to the capsid particles, changes in the brightness parameter are large (often exceeding an order of magnitude) and can readily be identified in fluorescence traces with an acquisition time of 10 s. Addition of inhibitors that either bind to the analyte (e.g. the CypA inhibitor cyclosporin A) or occupy the same binding site on capsid as the analyte (e.g. hexacarboxybenzene that binds to the same electropositive feature on capsid as dATP) reverse the fluorescence spikes and return the traces to background levels (Fig 2B, bottom). These intensity fluctuations can be quantified by calculating the brightness parameter (*B*), defined as the ratio of the variance (*σ*^2^) over the mean intensity (*μ*): *B* = *σ*^2^/*μ*.^36^ These calculations were performed using TRISTAN, a custom trace analysis software. Fig 2C shows the brightness parameters extracted from traces exemplified by those in Fig 2B. The brightness parameter of a monomeric analyte depends on the instrument (e.g. detection efficiency) and on the photophysical properties of the fluorophore (e.g. the brightness value of AF488 is larger than that of fluorescein). A large increase in brightness was observed in the presence of capsid particles, which was reversed upon addition of the corresponding inhibitor. We note that differences in the magnitude of the brightness parameter between different analytes cannot necessarily be interpreted with respect to relative affinities. As such, the primary use of this approach is to identify capsid-binding molecules rather than to measure binding affinities.

**Figure 2.**
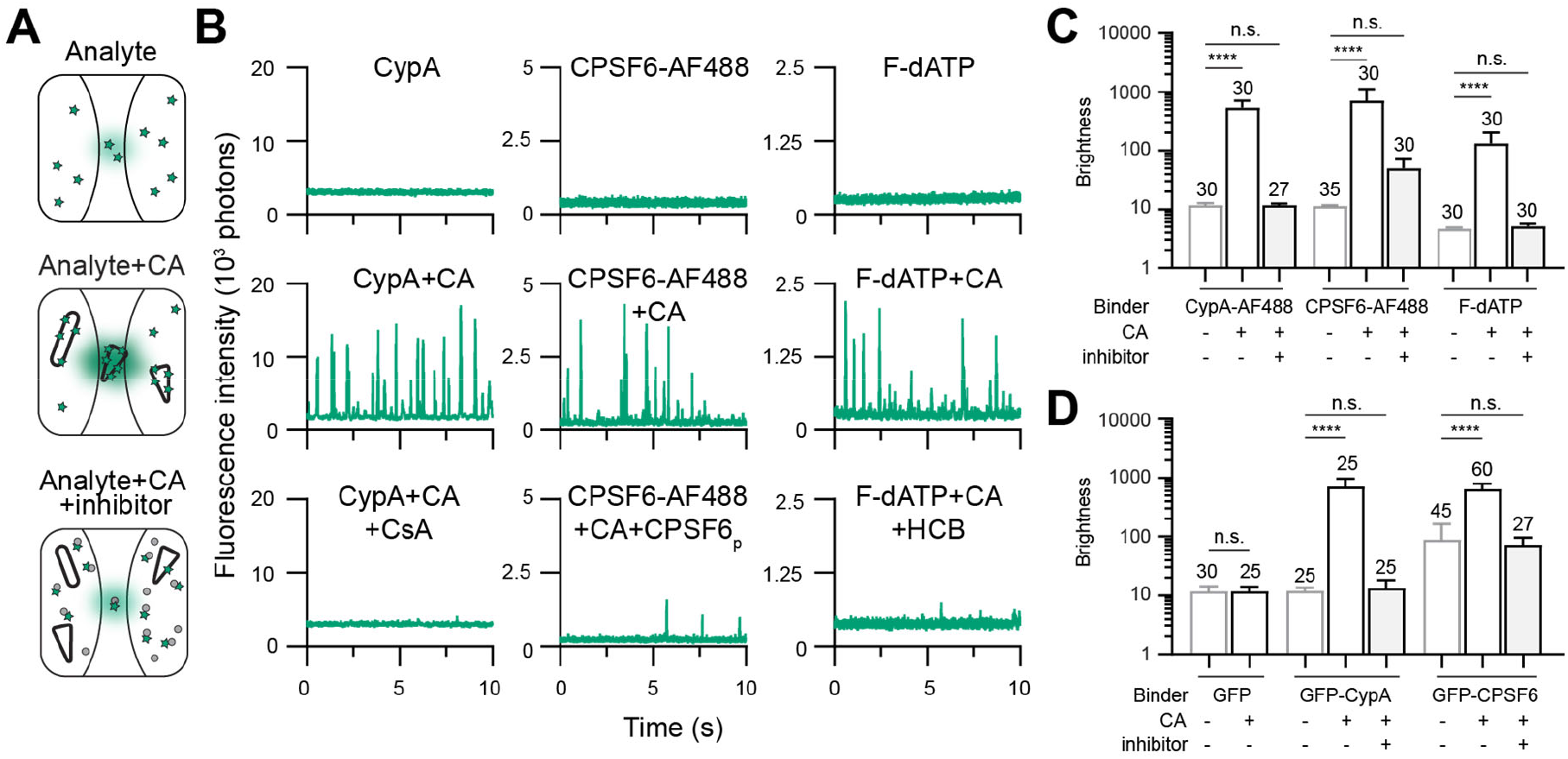
Detection of analyte binding to capsids via fluorescence spikes in single color assays. **A.** Confocal volume schematic of samples containing only analyte molecules (top) or mixtures of analyte and CA cones/tubes in the absence (middle) or presence of inhibitors (bottom). Inhibitors can either bind to the analyte (shown here) or to the interaction interfaces located on the CA lattice. **B.** Representative fluorescence traces of chemically labelled analytes (top), with capsid (middle) and with capsid+inhibitor (bottom): *left*, CypA-AF488 (∓CsA); *middle*, CPSF6-AF488 (∓unlabelled CPSF6 peptide); *right*, dATP labelled with fluorescein (∓HCB). **C./D.** Bar graphs of brightness values for chemically labelled analytes (C) or capsid-binding proteins fused to GFP expressed from *Leishmania* extract (D). The number of traces from ≥3 (C) or ≥2 (D) independent experiments are shown above the bars. Error bars represent the standard deviation. Comparisons using ordinary one-way ANOVA or unpaired t-test (GFP control), *p*≤0.0001 (****), *p*>0.05 (n.s.).

Next, we tested whether brightness analysis was compatible with different methods for analyte preparation and labelling. Cell-free expression systems provide a rapid and convenient approach for preparing proteins that are difficult to produce by other means. Here we used a *Leishmania* cell-free protein expression^31,32^ system to produce GFP, GFP-CypA, and full-length GFP-CPSF6 (Supporting Fig S1A) and examined the interactions of these proteins with capsid in the Brightness assay (Fig 2D, see Fig S1B for representative traces). The advantage of this system is that it allows rapid production of protein analytes (within ~2.5 h) in a microwell format in sufficient quantities for use in the assay without further purification. Both free GFP and GFP-CypA diffused as monomers in the absence of capsid. In the presence of capsid, free GFP remained monomeric whereas GFP-CypA showed a 1000-fold increase in brightness which was reversed by the addition of the inhibitor cyclosporin A. The intensity trace of GFP-CPSF6 without capsid showed that the full-length N-terminally tagged protein formed oligomeric species, yielding a 4–8-fold increase in the brightness parameter compared to monomeric GFP. In the presence of capsid, the brightness parameter further increased by about 10-fold, which was reversed by addition of unlabelled CPSF6 peptide that is sufficient to occupy the binding site on capsid. These observations also suggest that the oligomeric species interacts with the capsid via the canonical site. Indeed, oligomerization of CPSF6 may contribute to its physiological role in HIV infection by increasing avidity and mediating capsid disassembly.^23,38^

Finally, we synthesized a tris-NTA derivative of AF488 to label recombinant proteins with hexahistidine tag via formation of a non-covalent complex (Supporting Fig 3A) with a half-life of about 30 min.^39^ His-tagged CypA labelled with this compound showed the expected inhibitor-reversible increase in brightness in the presence of capsid while no increase was observed for a tris-NTA-AF488-labelled control protein of bacterial origin (LcrV) (Supporting Fig 3B). Taken together, our observations suggest that brightness analysis of capsid-binding analytes can be used with sample preparation and labelling methods that facilitate screening of potential new capsid-binding proteins.

**Figure 3.**
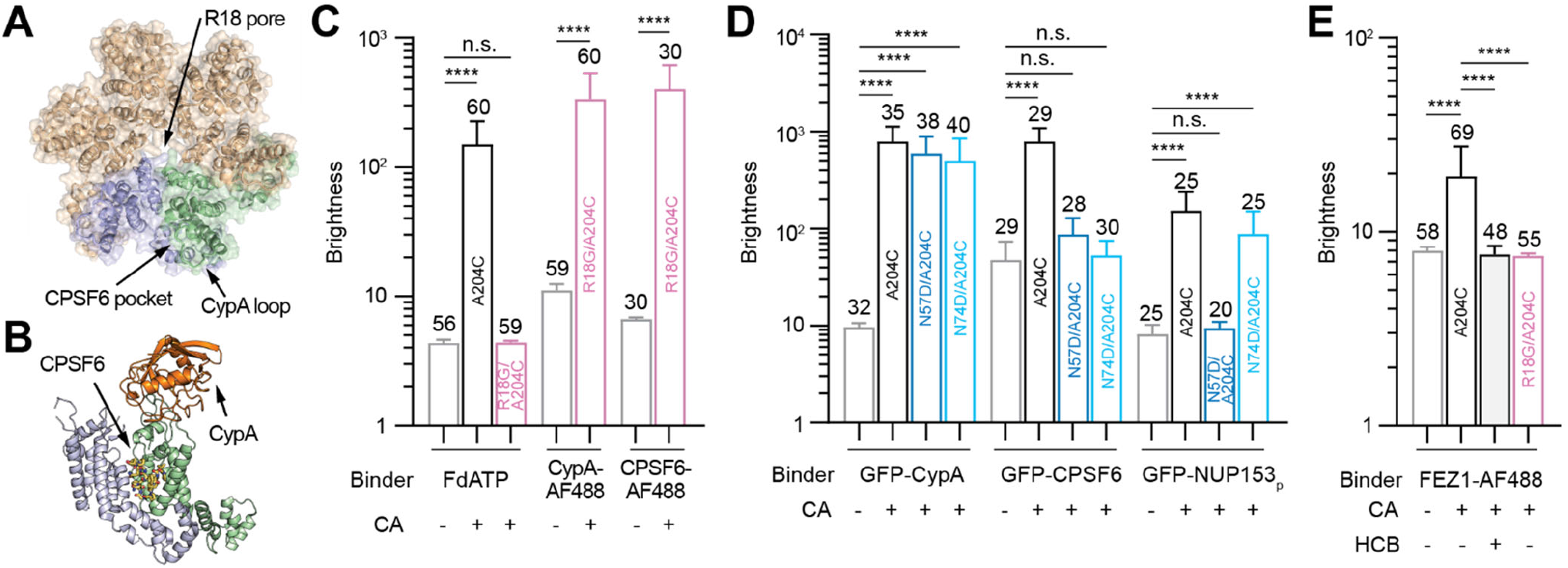
Identification of cofactor binding interfaces using CA mutants. **A.** Top view of a CA hexamer with two neighboring CA subunits shown in green and blue. Arrows point to cofactor binding interfaces. **B.** Side view of two neighboring CA subunits (green and blue) with bound CypA (orange) and CPSF6 peptide (stick representation). **C.** CA pore mutant R18G abolishes binding of fluorescent dATP but not of CypA or CPSF6 peptide. Bar graph of brightness values of F-dATP (50 nM), CypA-AF488 (50 nM) and CPSF6-AF488 (250 nM), in the absence of capsid (gray) or presence of self-assembled CA A204C (black) or R18G/A204C (violet). Traces were combined from at least 3 independent experiments. **D.** Bar graph of brightness values of GFP-CypA, GFP-CPSF6, GFP-Nup153p binding to self-assembled CA A204C (black), N57D/A204C (dark blue) and N74D/A204C (light blue) or in the absence of capsid (grey). B-values were obtained from cell-free expressed proteins from 2 different days. Error bars represent standard deviation. The number of B-values are displayed on top of each bar. **E.** FEZ1 peptide binds to the R18 pore. Bar graph of brightness values of FEZ1 peptide labelled with AF488 in the absence of capsid (grey) or presence of self-assembled CA A204C (black) HCB or R18G/A204C (violet). Statistics used ordinary one-way ANOVA or unpaired t-test, *p*≤0.0001 (****), *p*>0.05 (n.s.).

### Identification of analyte binding interfaces on capsid using binding site mutants

The HIV capsid comprises distinct binding sites for host proteins (Fig 3A and B), including: (1) The cyclophilin binding loop exposed on the external surface of each CA subunit binds CypA in the cytoplasm^40^ and the nuclear pore protein NUP358 upon arrival at the nucleus.^13^ (2) The CPSF6 binding pocket formed between the N-terminal and C-terminal domains of adjacent CA subunits in a hexamer binds phenylalanine-glycine containing peptides from nuclear protein CPSF6 as well as the nuclear pore protein NUP153.^14^ (3) The electropositive arginine ring formed at the center of each CA hexamer binds the adapter protein FEZ1 required for cytoplasmic capsid transport^7,41,42^ as well as small polyanionic molecules including nucleotides,^3^ inositol hexakisphosphate^2,43^ and the inhibitor hexacarboxybenzene.^3^

We took advantage of known amino acid substitutions in CA that selectively abolish binding of host molecules at either the central pore (R18G)^3^ or the CPSF6 binding pocket (N57D or N74D).^14^ Capsid particles assembled with CA A204C and either of these substitutions formed a mixture of capped tubes and cones (Supporting Fig S2). As expected, brightness analysis using capsid particles lacking the electropositive R18 ring showed that binding of F-dATP was abolished while CypA-AF488 and CPSF6-AF488 binding was unperturbed (Fig 3C). Similarly, binding of GFP-CPSF6 and the minimal peptide (residues 1407-1423) for capsid binding from NUP358 fused to GFP (GFP-NUP153p) was evident from the >10-fold increase in brightness in the presence of CA A204C particles (Fig 3D). This increase was ablated by substitution of residues forming critical hydrogen bonds with CPSF6 (N57 and N74) and NUP153 (N57).^14^ The host protein FEZ1 serves as an adaptor to link incoming capsids to kinesin motors for microtubule-based transport, whereby a recent study suggested that the FEZ1-capsid interaction is mediated by an unstructured acidic region that binds to the R18 pore.^7^ To test this requirement for binding, we measured binding of the corresponding FEZ1 peptide (residues 178-188) labelled with AF488. We observed a statistically significant 2–3-fold increase in the brightness parameter in the presence of capsid particles, which could be abolished either with the pore-binding inhibitor hexacarboxybenzene or by introduction of R18G (Fig 3E), supporting that the acidic peptide is a determinant for FEZ1 binding to the pore.^7^ Taken together these observations suggest that brightness analysis combined with a panel of binding site substitutions on the capsid are a useful tool to identify interaction sites for new host factors.

### Two-color coincidence detection of fluorescent analytes and capsid particles allows quantification of binding levels

Next, we assessed whether analyte-capsid complex formation could be measured quantitatively via two-color coincidence detection (TCCD)^29^ using AF568-labelled capsid particles (Supplementary Fig S2) as bait. Fluorescence intensity traces recorded with CypA-AF488 as prey (Fig 4A) show CypA peaks coincided with capsid peaks (Fig 4B, top). Addition of the inhibitor CsA resulted in the disappearance of the CypA peaks (Fig 4B, bottom), confirming that detection of coincident peaks was dependent on binding to capsid. A heatmap plot of the analyte intensity as a function of the coincident capsid intensity generated using our custom analysis software TRISTAN showed a linear relationship, i.e. larger capsids with higher intensity carry a proportionately larger number of analytes (Fig 4C). We note that the bright feature (large number of data points) in the bottom left quadrant of the heatmap plot arises from the large number of background events in both signals. The intensity ratio given by the slope of a linear fit line provides a measure proportional to the analyte:CA ratio at binding equilibrium.

**Figure 4.**
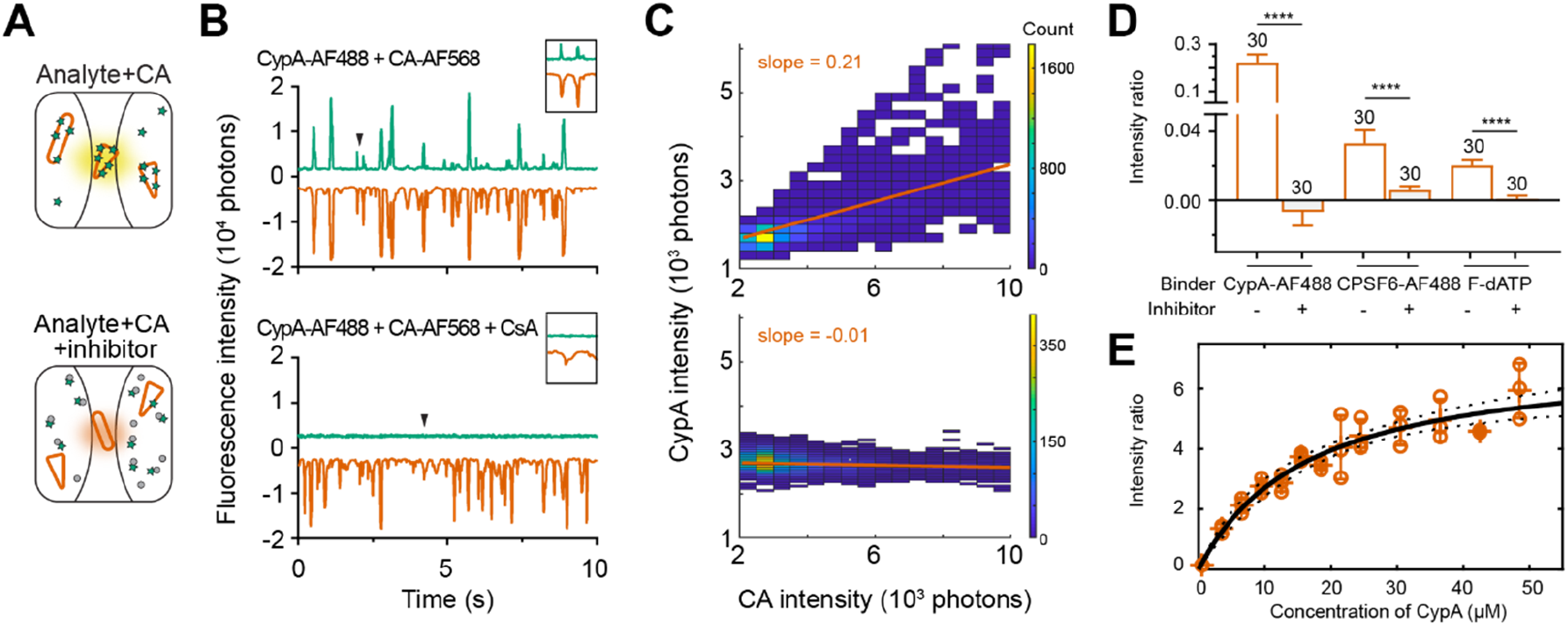
Detection of analyte binding to capsids via coincident peaks in two-color assays. **A.** Schematic of samples containing green-fluorescent analyte (represented as stars) and red-fluorescent CA cones/tubes in the absence (top) or presence (bottom) of inhibitors. **B.** Representative two-color fluorescence traces of CypA-AF488 (50 nM, green trace) binding to self-assembled CA A204C:CA K158C-AF568 (vermillion trace) in the absence (top) and presence (bottom) of the inhibitor CsA (5 μM). The insets show an enlarged view of the peaks indicated by the black arrowhead to reveal coincidence of peaks in the two channels. **C.** Heatmaps of the CypA-AF488 signal versus the CA K158C-AF568 signal from the traces shown in A. The slope of the line of best fit (vermillion) representing the CypA:CA signal intensity ratio was calculated using TRISTAN. **D.** Bar graph of intensity ratios representing binding (∓inhibitor) of AF488-labelled CypA (∓CsA), fluorescein-labelled dATP (∓HCB) and AF488-labelled CPSF6 peptide (∓unlabelled CPSF6 peptide). Error bars represent the standard deviation. The number of traces (combined from three independent experiments) is indicated above each bar. Comparisons using two-tailed unpaired t-test, p≤0.0001 (****). **E.** Equilibrium binding curve obtained by plotting the CypA:CA intensity ratio as a function of CypA concentration. The CA concentration was kept at 1 μM. Error bars represent the standard deviation of measurements from three repeats. The fit (black line) of an equilibrium binding model provided values for the dissociation constant (*K*D=16.5 μM [10.1−23 μM, 95% CIs]).

Using TCCD, we determined the intensity ratio for cofactors CypA-AF488, CPSF6-AF488 or F-dATP interacting with capsid added in a large molar excess to the binding reaction (Fig 4D, also see Supporting Fig S3C for hexahistidine-tagged CypA labelled with tris-NTA-AF488). The analyte:CA intensity ratio determined for CypA-AF488 was about 5-fold larger than for CPSF6-AF488, reflecting the differences in affinity (*K*D of ~10 μM for CypA ^16,22,23,40,44^ and ~50 μM for CPSF6^14^). The low intensity ratio determined for F-dATP reflects both the lower number of binding sites on the capsid (1 pore per CA hexamer) as well as the lower emission of photons by fluorescein compared to AF488. Addition of the corresponding inhibitor resulted in a drop of the intensity ratios to values close to zero, confirming inhibition of capsid binding. Examples for TCCD capsid binding analysis of GFP-tagged analytes, including oligomeric proteins which show characteristic features in the heatmap plots of an lyte intensity as a function of capsid intensity, are shown in Supporting Fig S4.

Finally, we used TCCD to measure an equilibrium binding curve for CypA-AF488 added at a range of concentrations (1–50 μM) to AF568-labelled capsid particles (equivalent CA concentration of 1 μM) (Fig 4E). A fit of an equilibrium binding model provided an estimate of 16.5 μM for the dissociation constant of the interaction, consistent with previous measurements using a range of different techniques.^16,22,23,40,44^ Taken together our observations show that the intensity ratio analysis of two-color data provides a simple approach requiring low amounts of reagents to quantify binding levels and affinities. Molar ratios could in principle be obtained by calibration of the fluorescence intensities of each fluorophore.

## CONCLUSION

Here we characterize two fluorescence fluctuation spectroscopy assays for measuring binding of fluorescent analytes to stabilized HIV-1 capsid particles: (1) Brightness analysis of fluorescent analyte traces reveals binding via accumulation of multiple analytes onto the lattice, leading to the appearance of large intensity spikes. This mode is primarily suited for initial identification of molecules that interact with capsid, (2) Intensity ratio analysis of coincident analyte and capsid signals in two-color experiments to quantitate the level of analyte binding at equilibrium. The simple format of the assay is facilitated by the availability of low-cost 3D-printed confocal spectroscopy systems.^45^

Advantages of the binding assays include short measurement times facilitated by the well-scanning system, a low reagent requirement and compatibility with complex mixtures, allowing preparation of protein analytes in a microplate format using cell-free expression systems. Measurements are performed on freely diffusing particles in solution, avoiding the need to troubleshoot methods for specific capture onto surfaces. The high number of binding sites on the capsid particles allows detection of analytes with high sensitivity. Applications include screening to identify new capsid-binding host cell proteins (especially when used in conjunction with cell-free expression), and dissection of capsid binding interfaces using a panel of CA mutants. In addition, two-color measurements enable quantitative measurement to compare the level of recruitment of different ligands or to determine affinity. This system could be expanded to include protocols for calibration of fluorophore intensities, which would allow the determination of molar analyte:CA ratios. Importantly, these methods are also applicable to study capsid-host interactions of other viruses where capsid plays a central role in infection. More generally, the approaches presented here could be extended to detect analyte binding to other types of multivalent scaffolds such as protein assemblies, DNA structures or liposomes, with the possibility of engineering polymeric or supramolecular interaction platforms to cluster analytes for interaction screening with high sensitivity.

## SUPPORTING INFORMATION

Supporting information (PDF) includes Supporting Materials and Methods, Supporting Figures (Cell free expression of host proteins fused to GFP; Ultrastructure of self-assembled CA A204C particles with amino acid substitutions in cofactor binding interfaces; Detection of hexahistidine-tagged proteins labelled with tris-NTA-AF488 dyes binding to AF568-labelled capsids; Interpretation of TCCD data with monomeric and oligomeric analytes) and Supporting Discussion. This material is available free of charge via the Internet at http://pubs.acs.org.

## Author Contributions

DL - protein production/labelling, CA assembly, EM, assay development, data acquisition/analysis, original draft of the manuscript with TB; JW - data analysis software, advise on experimental setup; CD - protein production, assay development and acquisition of CPSF6 binding data; AT, JS - assay development and analysis; DH - production of *Leishmania* extract; AB - optimisation of cell-free protein expression; VS - protein production, supervision, advise on CA assembly and HIV cofactor usage; ST - supervision, advise on HIV cofactor usage; ES, YG - supervision of cell-free expression and assay development, confocal microscope setup; TB, DJ - conceptualization, supervision. All authors revised the manuscript.

## FUNDING AND ACKNOWLEDGMENT

This work was supported by the National Health and Medical Research Council of Australia (APP1098870, APP1182212, APP1110116). DL received an Australian Government Research Training Program Scholarship. We thank Stephanie Xu for providing LcrV; James Brown for discussions; Wang Peng for help with EM. The authors acknowledge the instruments and scientific and technical assistance of Microscopy Australia at the Electron Microscope Unit, UNSW, a facility that is funded by the University, and State and Federal Governments.

## ABBREVIATIONS

CA: capsid protein
CPSF6: cleavage and polyadenylation specificity factor subunit 6
CsA: cyclosporin A
CypA: cyclophilin A
F-dATP: fluorescein labelled dATP
FEZ1: fasciculation and elongation protein 1
HCB: hexacarboxybenzene
IP6: inositol hexakisphosphate
MVP: major vault protein
NUP153: nucleoporin 153
TCCD: two-color coincidence detection
TRISTAN: Two Reagents Incident Spectroscopic Analysis
VIM: vimentin.

## Table of Contents artwork

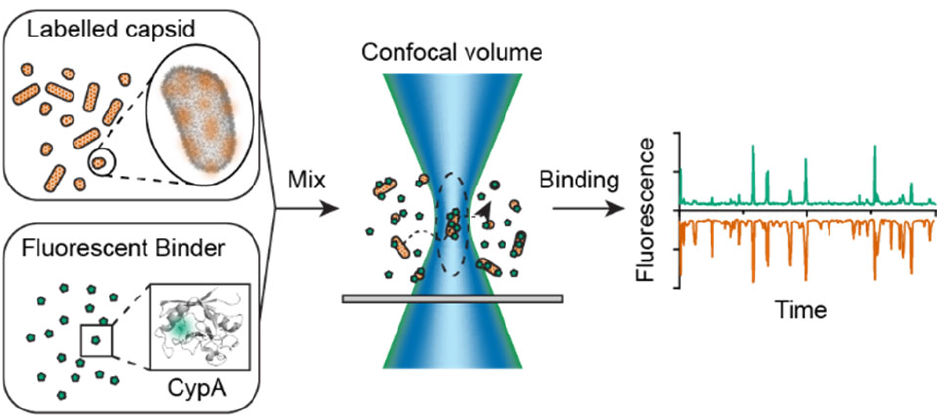

