## Supplementary for "Rapid HIV-1 capsid interaction screening using fluorescence fluctuation spectroscopy"

### Supporting Methods

**Purification of recombinant HIV-1 CA.** HIV-1 CA protein was expressed from a pET11a construct. Point mutations (K158C, A204C, R18G/A204C, N57D/A204C and N74D/A204C) were introduced using site directed mutagenesis. Protein was expressed from the *Escherichia coli* strain Rosetta2 (DE3, pLysS). Cell pellets were resuspended in lysis buffer (50 mM Tris, pH 8; 2 mM DTT; 0.02% w/v $NaN_3$) and lysed by sonication. The clarified lysate was incubated with 2.5 M NaCl to induce CA assembly. The solution was centrifuged, and the pellet containing assembled CA was redissolved in lysis buffer. NaCl fractionation was repeated once more and the soluble capsid was subjected to subtractive anion exchange chromatography using a 5 mL HiTrap Q HP column (GE Healthcare, 17115301). CA was collected in the flowthrough and concentrated to >100 µM (2.5 mg/mL). Protein concentration was determined via the Bradford assay (Thermo Fisher Scientific, 23236) using BSA as standard. Aliquots of 20 µL were snap-frozen in liquid nitrogen and stored at -80 °C.

**Purification of recombinant $His_6$-tagged proteins.** A construct producing CypA with an N-terminal hexahistidine tag ($His_6$-CypA), from the pET-MCSIII vector, was a kind gift from Nicholas Dixon. Expression was performed in Rosetta2 (DE3, pLysS) cells. Cells were grown at 37 °C in Luria Broth supplemented with ampicillin (100 µg/mL) and chloramphenicol (34 µg/mL) to $OD_{600\ nm}$ = 0.5 and induced with 1 mM IPTG for 3 h at 37°C. Cells were resuspended in 50 mM sodium phosphate, pH 6.5; 300 mM NaCl; 10 mM imidazole supplemented with 1× cOmplete EDTA-free protease inhibitor (Roche, Switzerland) and lysed by sonication. Lysate was incubated with Ni-NTA resin (2 mL, 50% slurry, Qiagen) for 1 hour at 4 °C and washed with lysis buffer containing 20 mM imidazole. Bound $His_6$-CypA was eluted with a lysis buffer containing 250 mM imidazole and dialysed into 25 mM sodium phosphate, pH 6.5; 1 mM DTT; 0.02% w/v $NaN_3$. Finally, size exclusion chromatography was performed over a Superdex 200 Increase 10/300 GL column (GE Healthcare, 17517501) equilibrated in 25 mM sodium phosphate, pH 6.5; 150 mM NaCl, 1 mM DTT, 0.02% w/v $NaN_3$. The purified $His_6$-CypA was concentrated to ~100 µM and snap-frozen in liquid nitrogen for storage at -40 °C. Purified, recombinant LcrV (carrying a mutation, C293T, and a 6xHis tag on the N-terminus) from the type III secretion system of *Yersinia pestis* was gifted by Stephanie Xu.

**Synthesis of tris-NTA Alexa Fluor 488 (tris-NTA-AF488).** Alexa Fluor 488 sulfodichlorophenol ester (1 mg, 1.2 µmol, Thermo Fisher Scientific, A30052) and tris-NTA trifluoroacetic acid salt (2 mg, 1.2 µmol, Toronto Research Chemicals, N925005) were dissolved in dry dimethyl sulfoxide (48 µL). N,N-diisopropylethylamine (5 µL) was added and the solution was mixed. The progress of the reaction was monitored by thin layer chromatography on silica gel plates developed in methanol:water (4:1, v/v). After a reaction time of 48 h at room temperature in the dark, the mixture was concentrated under reduced pressure and the residue was taken up in water (400 µL). The desired product was then purified by HPLC using an Alltima HP C18 column (Alltech) operated at a flow rate 1 mL/min with mixtures of solvent A (water containing 0.1% trifluoroacetic acid) and solvent B (acetonitrile containing 0.1% trifluoroacetic acid). After injection of product mixture, compounds were eluted with 12.5% B for 10 min, whereby tris-NTA-AF488 eluted at 9 min. The column was then washed with 25% solvent B (5 min) and re-equilibrated with 12.5% solvent B (8 min). The fractions containing tris-NTA-AF488 were combined and evaporated to dryness under reduced pressure to yield an orange solid. The overall yield of the reaction was 62%. The purity of the final product was confirmed by thin layer chromatography and HPLC. The absorption maximum of the final product in water was 495 nm.

**Production of cell-free lysate.** *Leishmania tarentolae* Parrot strain was obtained as LEXSY host P10 from Jena Bioscience GmbH, Jena, Germany and cultured in TBGG medium containing 0.2% v/v Penicillin/Streptomycin (Life Technologies) and 0.05% w/v Hemin (MP Biomedical). Cells were harvested by centrifugation at 2500 g, washed twice by resuspension in 45 mM HEPES, pH 7.6, containing 250 mM sucrose, 100 mM potassium acetate and 3 mM magnesium acetate and resuspended to a density of 0.25 g cells/g suspension. Cells were placed in a cell disruption vessel (Parr Instruments, USA) and incubated under 7000 kPa nitrogen for 45 minutes, then lysed by rapid release of pressure. The lysate was clarified by sequential centrifugation at 10000 g and 30000 g and anti-splice leader oligonucleotide was added to 10 M. The lysate was then desalted into 45 mM HEPES, pH 7.6, containing 100 mM potassium acetate and 3 mM magnesium acetate and snap-frozen until required.

Cell-free lysate was supplemented with a feeding solution containing nucleotides, amino acids, T7 polymerase, HEPES buffer, and a creatine/creatine kinase ATP regeneration system at a ratio of lysate to feed solution of 0.21, and a final $Mg^{2+}$ concentration of 6 mM. Purified plasmid DNA, at a concentration of 60 nM, was added to the expression reaction at a ratio of 1:9 (v/v), and the reaction was allowed to proceed for 2.5 h at 27°C. Fluorescently tagged/labelled expressed protein was detected both before and after SDS-PAGE using a Chemidoc MP imaging system (Bio-Rad, Laboratories Pty. Ltd.,Gladesville, NSW, Australia).

**Supporting Figures**

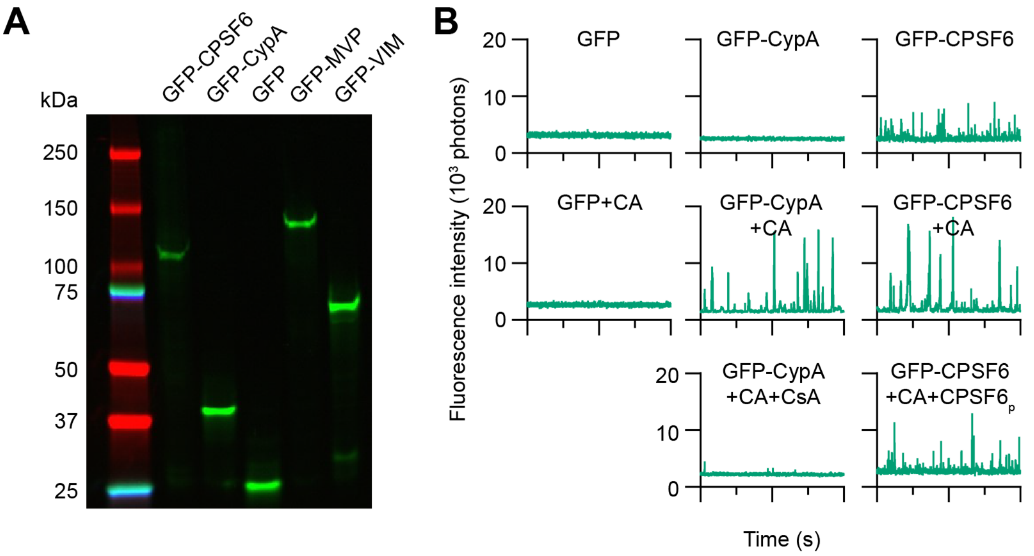


**Figure S1. Cell-free expression of host proteins fused to GFP. A.** SDS-PAGE analysis with fluorescence detection of GFP-CPSF6, GFP-CypA, GFP, GFP-MVP and GFP-VIM expressed using *Leishmania* extract. **B.** Representative fluorescence fluctuation spectroscopy traces of capsid-binding proteins fused to GFP produced by cell-free expression (top) and or mixtures of these proteins with capsid particles in the absence (middle) or presence (bottom) of inhibitor: GFP only (control), GFP-CypA (∓CsA), GFP-CPSF6 (∓unlabelled CPSF6 peptide).

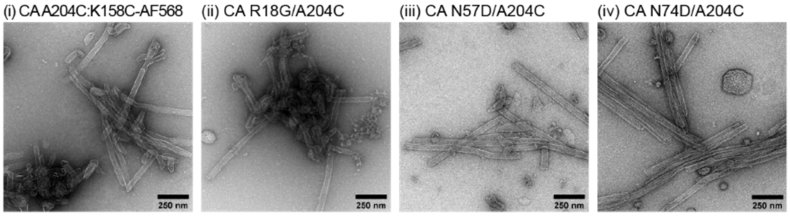


**Figure S2. Ultrastructure of self-assembled CA A204C particles with amino acid substitutions in cofactor binding interfaces.** Negative staining electron micrographs of disulfide cross-linked structures formed by self-assembly of (i) a mixture of CA A204C and CA K158C-AF568 (76 µM:4 µM), (ii) CA R18G/A204C (80 µM), (iii) CA N57D/A204C (80 µM), (iv) CA N74D/A204C (80 µM).

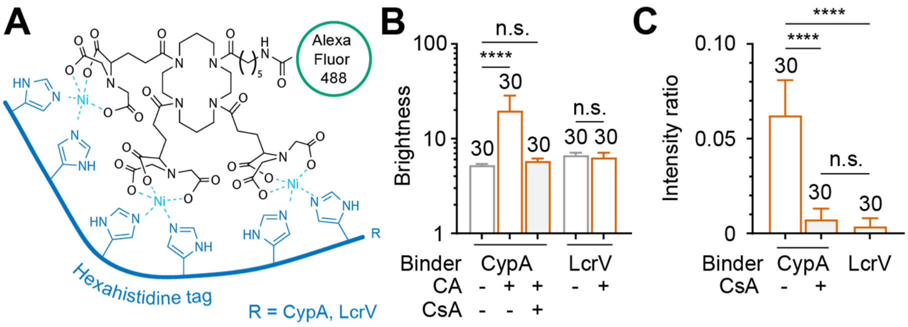


**Figure S3. Detection of hexahistidine-tagged proteins labelled with tris-NTA-AF488 dyes binding to AF568-labelled capsids. A.** Labelling scheme. **B.** Bar graph of brightness values of CypA (∓cyclosporin A, CsA) and LcrV (negative control) labelled with tris-NTA-AF488 in the absence and presence of CA cones/tubes. The number of traces (combined from three independent experiments) is indicated above each bar. **C.** Bar graph of analyte:CA intensity ratios obtained by two-colour coincidence detection for CypA (∓cyclosporin A, CsA) and LcrV (negative control) labelled with tris-NTA-AF488**.** The number of traces (combined from three independent experiments) is indicated above each bar. Error bars represent the standard deviation of the mean. Comparisons using ordinary one-way ANOVA (CypA) or unpaired t-test (LcrV), p≤0.0001 (****), p>0.05 (n.s.).

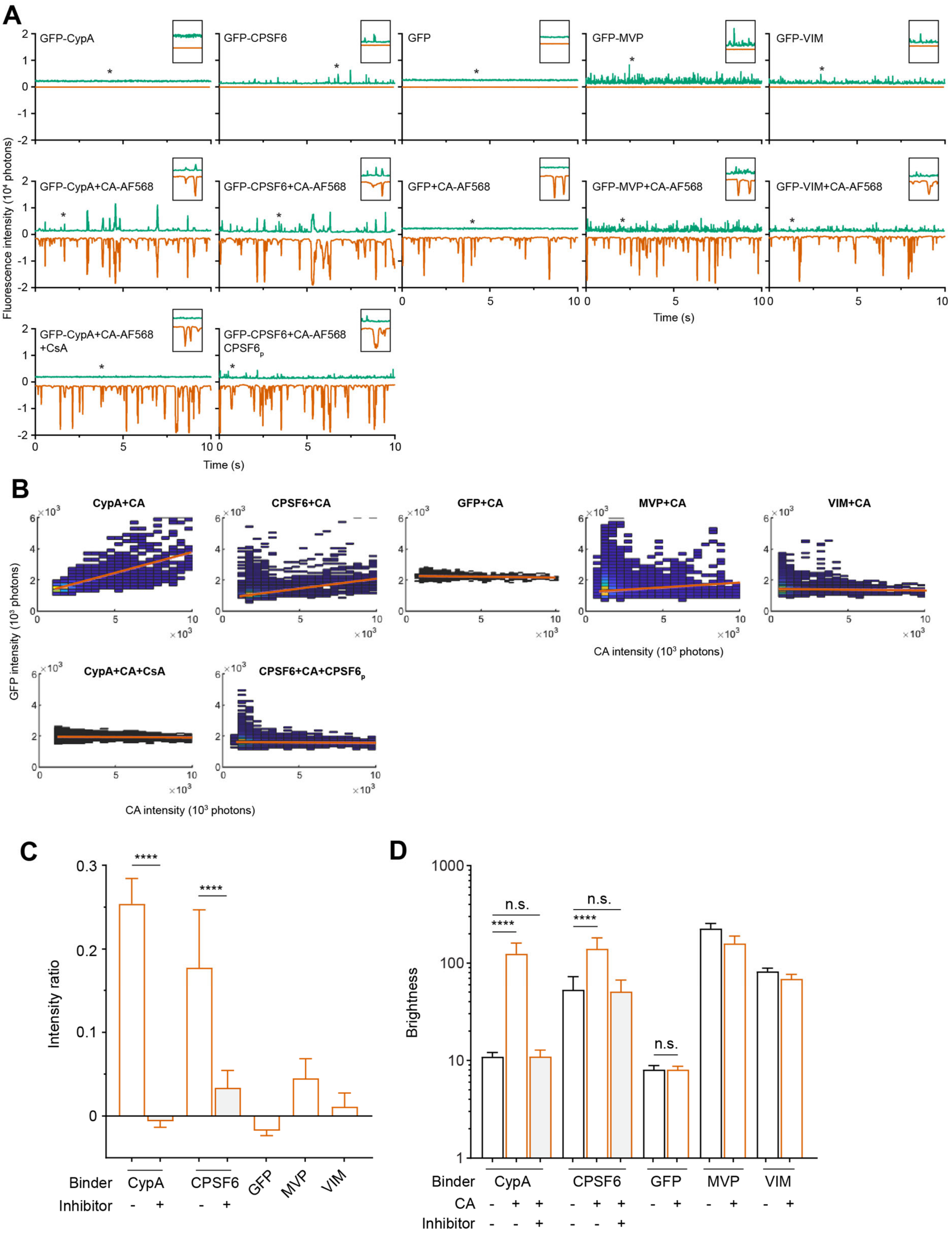

**Figure S4. Interpretation of two-colour data with monomeric and oligomeric analytes. A.** Representative two-colour fluorescence traces of GFP-tagged proteins (green trace) produced by cell-free expression and self-assembled CA A204C:CA K158C-AF568 particles (vermillion trace) comparing the following conditions: *top row*, GFP-tagged proteins without capsid; *middle row*, GFP-tagged proteins in the presence of AF568-labelled capsids; *bottom row*, GFP-tagged proteins in the presence of AF568-labelled capsids and corresponding inhibitor. The insets show an enlarged view of the peaks indicated by the asterisk. Analytes include (from left to right): GFP-CypA ∓cyclosporin A (CsA) [monomeric binder]; GFP-CPSF6 ∓unlabelled CPSF6 peptide [oligomeric binder]; GFP [monomeric non-binding control]; GFP-tagged major vault protein (GFP-MVP) [oligomeric]; GFP-tagged vimentin (GFP-VIM) [oligomeric]. **B.** Heatmaps of the GFP-analyte signal versus the CA K158C-AF568 signal from the traces shown in A (middle and bottom rows). The slope of the line of best fit (vermillion) representing the analyte:CA signal intensity ratio was calculated using TRISTAN. **C.** Bar graph of intensity ratios. Error bars represent the standard deviation. Comparisons using two-tailed unpaired t-test, $p \leq 0.0001$ (****). **D.** Bar graphs of brightness values calculated from traces of the GFP channel (single-colour analysis of the data shown in A). Comparisons using ordinary one-way ANOVA or two-tailed unpaired t-test, $p \leq 0.0001$ (****), $p > 0.05$ (n.s.).

### Supporting discussion

**Analysis of monomeric and oligomeric analytes by TCCD.** Monomeric analytes such as GFP-CypA or GFP in the absence of capsid show flat traces (Fig S4A, top). Addition of labelled capsid to capsid-binding analytes results in the appearance of intensity peaks in the analyte channel that coincide with the capsid peaks (GFP-CypA; Fig S4A, middle). The heatmap generated from the traces of a monomeric binder shows a linear dependence with a positive slope when the analyte signal is plotted against the corresponding capsid signal (GFP-CypA; Fig S4B and C). These coincident peaks are not observed when binding is inhibited (GFP-CypA+CsA; Fig S4A, bottom) or absent (GFP; Fig S4A, middle). The corresponding heatmaps for non-binding monomeric analytes are characterised by a line of best fit with a slope close to zero (GFP-CypA+CsA or GFP; Fig S4B and C). Brightness analysis of the analyte traces can be used to confirm the two-colour analysis described above, showing an increase in brightness upon addition of capsid only for analytes that bind to capsid (Fig S4D).

Binding to capsid can be difficult to ascertain for oligomeric analytes by inspecting traces from two-colour measurements. Example traces for GFP-CPSF6 exhibit peaks in the absence of capsid (Fig S4A, top) indicating that the protein forms oligomeric species. Binding of GFP-CPSF6 oligomers to capsid particles is apparent from the coincident peaks in both channels, whereas GFP-CPSF6 oligomers that remain unbound do not have coincident capsid peaks. The corresponding heatmap analysis (Fig S4B) shows a linear relationship with a positive slope (arising from GFP-CPSF6 oligomers in complex with capsid) as well as intensity along the y-axis (arising from unbound GFP-CPSF6 oligomers). Accumulation of analyte onto the capsid can also be confirmed by brightness analysis of the analyte traces (Fig S4D). When CPSF6 binding is outcompeted by the addition of unlabelled CPSF6 peptide, the heatmap has intensity along the x- and y-axes, giving a characteristic L shape (Fig S4B).

Two-colour analysis of GFP-tagged major vault protein (GFP-MVP) and GFP-tagged vimentin (GFP-VIM) give rise to primarily L-shaped heatmaps (Fig S4C). The line of best fit for GFP-VIM has a slope close to zero indicating the absence of complex formation with capsid. The line of best fit in the heatmap for GFP-MVP shows a small positive slope, which is presumably due to coincident peaks that arise by chance rather than by capsid binding (note the high frequency of peaks in the corresponding GFP-MVP traces shown in Fig S4A, which could be alleviated by dilution of the analyte). Overall, the analyte:CA intensity ratio of GFP-MVP and GFP-VIM show little to no binding to capsid (Fig S4B/C) and this conclusion is further supported by brightness analysis of the analyte traces (Fig S4D).
